# Type I Interferon production in myeloid cells is regulated by factors independent of *Ptpn22*

**DOI:** 10.1101/2025.03.20.644450

**Authors:** Jenna R. Barnes, Anam Fatima Shaikh, Alec M. Bevis, Tammy R. Cockerham, Robin C. Orozco

## Abstract

The immune regulatory gene *PTPN22* is expressed in all immune cells and encodes Lyp in humans and the ortholog PEP in mice. The *PTPN22* alternative allele, 1858C>T, is expressed in 5-15% of the North American population and is strongly associated with the development of autoimmune disease while simultaneously capable of providing protection during virus infection and cancer. In murine models, significant progress has been made in elucidating the molecular mechanisms that PEP and its pro-autoimmune variant (PEP-R619W) modulate T cell function, yet their influence on non-T cell pathways, such as antigen presenting cell cytokine production, remains less defined. Previously, it was reported that PEP promotes type I interferon (IFN-I) production in dendritic cells (DCs) and macrophages following TLR4 stimulus. Here, we show that contrary to previous results, both PEP-WT and the PEP-R619W variant do not promote IFN-I production in DCs and macrophages following exposure to LPS, 3p-hpRNA, or coronavirus MHV A59. We attribute the prior findings to mouse strain-specific differences and conclude that factors independent of PEP may be regulating IFN-I production in these studies. We further show that PEP and its R619W variant distinctly modulate the production of TNFα, IL-12 and IL-2 in DCs following LPS stimulus. Taken together, our results challenge the current understanding of the role of PEP during inflammation while providing new insight into how the PEP-R619W variant may alter myeloid cell function during disease.

## INTRODUCTION

Allelic variation in immune-regulatory genes can significantly impact disease outcomes ^1, 2^. One variant of interest is the 1858C>T allele of the Protein Tyrosine Phosphatase Non-Receptor Type 22 gene (*PTPN22*). *PTPN22* is constitutively expressed in all immune cells and encodes for the phosphatase lymphoid protein (Lyp), and its murine ortholog, PEST-domain enriched phosphatase (PEP). The *PTPN22* 1858C>T allele causes an amino acid substitution of arginine (R) to tryptophan (W) at position 620 in humans (Lyp-R620W). It is expressed in 5-15% of the North American population and is considered the highest non-HLA risk allele for the development of autoimmunity, including type I diabetes, rheumatoid arthritis, and systemic lupus erythematous^3-11^. To investigate the role of *PTPN22* and its autoimmunity-associated 1858C>T variant during inflammation, researchers often employ mouse models that lack expression of *Ptpn22* (PEP-null). However, the LYP-R620W variant, or the murine equivalent PEP-R619W variant, does not cause a loss of protein expression, but rather altered function^12, 13^. Thus, using PEP-null models to investigate how the *PTPN22* 1858C>T autoimmunity-associated allele drives inflammation may not always be appropriate as PEP-null and PEP-R619W mice can have differing immune phenotypes at homeostasis and during disease ^14-20^.

Previously, it was reported that PEP promotes type I interferon (IFN-I) production in dendritic cells (DCs) and macrophages following stimulus with bacterial lipopolysaccharide (LPS) or Poly(I:C)), Toll Like Receptor 4 (TLR4) and Toll Like Receptor 3 (TLR3) agonists, respectively ^21^. These studies were performed using PEP-null mice available at The Jackson Laboratory (PEP-null JAX: strain number #028977) compared to C57BL/6J wild-type mice as the recommended controls per The Jackson Laboratory at that time. However, after these studies were published, the Jackson Laboratory discovered that the PEP-null JAX strain contain multiple single nucleotide polymorphisms (SNPs). These SNPs indicate the PEP-null JAX strain is on a mixed C57BL/6J;C57BL/6N background, despite reported backcrossing to the C57BL/6J strain ^22^. This is significant because the C57BL/6N SNPs present in this mouse model may impact immune responses such as IFN-I production. This poses concern that our current understanding of *Ptpn22* and how it regulates inflammation may have been confounded by mouse strain-specific genetic variation.

To enable a more rigorous investigation into defining how PEP and the PEP-R619W variant, regulate inflammation and control for potential C57BL/6N-related genetic variation that could confound our results, our research team employs C57BL/6J CRISPR-Cas9-generated PEP-null and PEP-R619W mouse models, which have been maintained on a C57BL/6J background. Here, we employ these genetic models to determine if previous data regarding the impact of PEP on IFN-I production could be recapitulated and define if PEP and the PEP-R619W variant impacted the production of other cytokines in dendritic cells (DCs). We report that neither PEP nor the PEP-R619W variant regulates IFNβ production in DCs or macrophages following LPS or viral RNA stimulation. However, we found that both PEP and PEP-R619W differentially modulate the production of other pro-inflammatory cytokines by dendritic cells, including IL-2, IL-12, and TNFα.

## RESULTS

### Strain-specific differences impact IFNβ production in PEP-null bone marrow dendritic cells (BMDCs)

Previously, it was shown that PEP-null bone marrow-derived dendritic cells (BMDCs) and bone marrow macrophages (BMMs) produce reduced amounts of IFN-I when stimulated with bacterial lipopolysaccharide (LPS) or Poly(I:C)), TLR4 and TLR3 agonists respectively ^21^. However, the PEP-null JAX strain was discovered to have multiple SNPs associated with a mixed genetic background, posing the concern that our current understanding of the role of PEP in regulating IFN-I production may have been influenced by mouse strain-specific differences, such as genetic variation. Using our C57BL/6J CRISPR-Cas9-generated PEP-null model (PEP-null CRISPR), we first set out to determine if previous data regarding the impact of PEP on IFN-I production could be recapitulated. Using qRT-PCR, we confirmed that our PEP-null CRISPR mice do not express quantifiable amounts of *Ptpn22* mRNA (Figure 1A). Then, using FLT3-ligand differentiated BMDCs from C57BL/6J PEP-WT, PEP-null CRISPR, and PEP-null JAX mice we measured for IFNβ production following LPS stimulation. While IFNβ concentration was comparable between PEP-WT and our PEP-null CRISPR mice, BMDCs from the commercial PEP-null JAX strain showed significantly less IFNβ production (Figure 1B). These data successfully recapitulate prior results, but through the use of our CRISPR-Cas9-generated PEP-null model, we observe that PEP does not promote IFNβ production. The differences between the PEP-null JAX and PEP-null CRISPR strains suggest that differences associated with the C57BL/6N background, rather than *Ptpn22* deficiency alone, may account for the previously reported phenotype.

**Figure 1.**
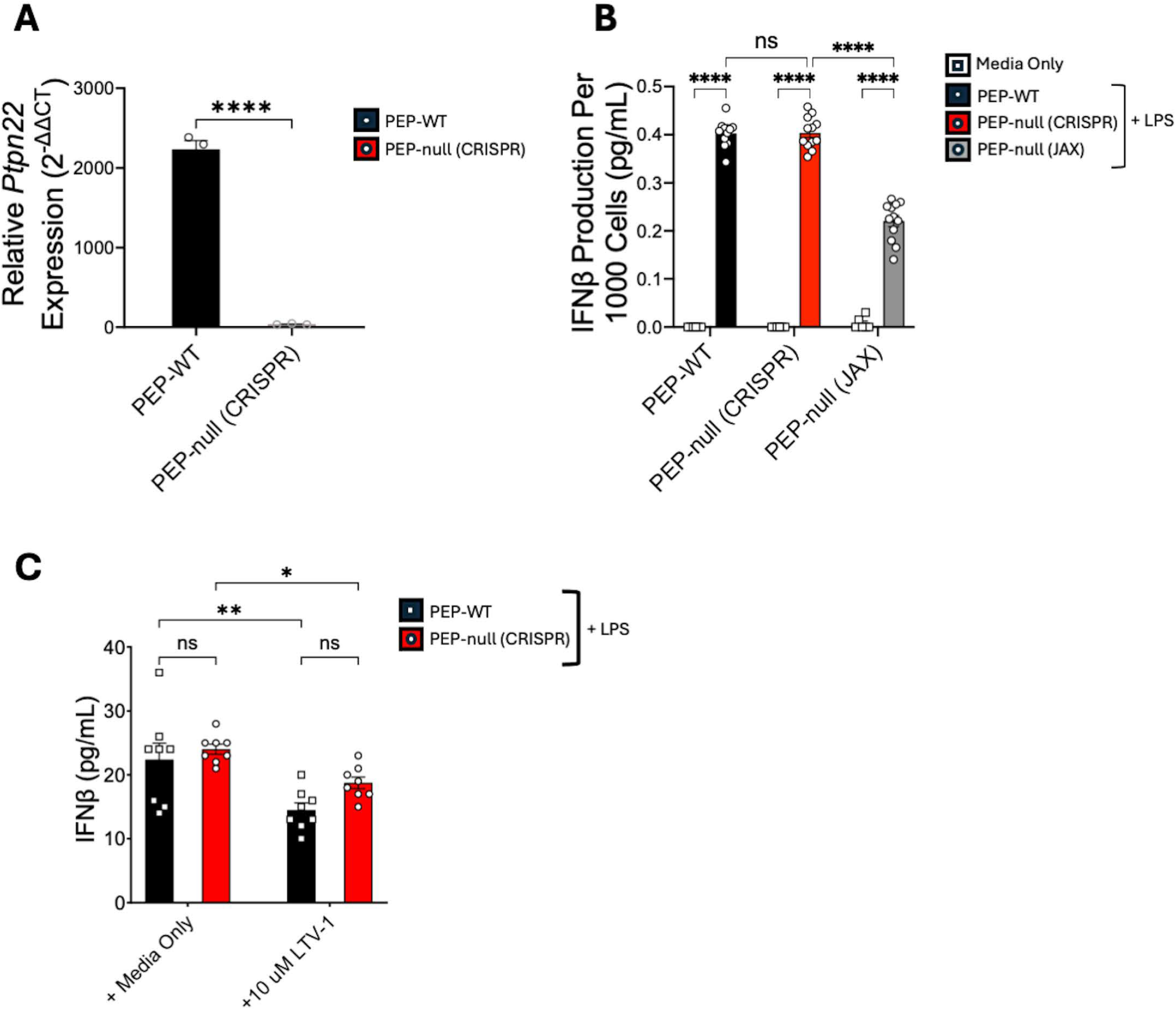
Strain-specific differences impact IFNβ production in PEP-null BMDCs. Gene expression values of *Ptpn22* relative to *Gapdh* were determined by qRT-PCR for naïve PEP-WT (black) and PEP-null (CRISPR) (red) splenocytes, each dot represents a mouse **(A)**. FLT3-L BMDCs were cultivated and differentiated from C57BL/6J PEP-WT (black), CRISPR/Cas9-generated PEP-null (red), and PEP-null mice from the Jackson Laboratory (JAX stock #028977) (gray). Cultures were incubated with LPS (500 ng/mL) for 18 hours. The supernatant was used to determine IFNβ concentration (ELISA) **(B)**. PEP-WT and PEP-null (CRISPR) BMMs were incubated with LPS (500 ng/mL) and 10 μM LTV-1 for 18 hours. The supernatant was used to determine IFNβ concentration (ELISA) **(C)**. ns= no significance; *p<0.01; **p<0.001; ****p<0.0001; Unpaired T Test (A), Two Way ANOVA with Tukey post hoc analysis (**B**,**C**). Data is pooled from 2 independent experiments. Each dot represents a biological replicate.

In addition to the PEP-null JAX model, studies have also utilized chemical inhibition of PEP to evaluate its role in IFN-I production and activity ^21, 23^. However, varying results regarding PEP inhibitors and their specificity have been reported ^24^. To evaluate the specificity of the PEP inhibitor LTV-1 during IFN-I production, we stimulated PEP-WT and PEP-null CRISPR bone marrow macrophages (BMMs) with LPS in the presence or absence of LTV-1 to inhibit PEP’s enzymatic activity. We observed reduced IFNβ production in both PEP-WT and PEP-null CRISPR BMMs treated with LTV-1, but no significant differences in IFNβ production between the two genotypes (Figure 1C). These data suggests that the reduction in IFNβ production is due to off-target effects of LTV-1, rather than inhibiting PEP. Given the potential confounding factors associated with the PEP-null JAX mice and off-target effects of LTV-1, we proceeded with our studies to define the impact of PEP and the PEP-R619W variant on inflammation with only our C57BL/6J CRISPR generated mouse strains.

### PEP and PEP-R619W do not mediate IFNβ production in myeloid cells

Our initial results showed comparable IFNβ production between PEP-WT and PEP-null CRISPR BMDCs following TLR4 stimulation via LPS. However, beyond TLRs, additional pattern recognition receptors (PRRs) such as RIG-I Like Receptors (RLRs) induce large quantities of IFN-I in response to virus infection ^25-27^. Further, it is unknown if the autoimmunity-associated PEP-R619W variant, which alters protein function rather than expression, uniquely affects IFN-I production compared to PEP-WT. To determine if the PEP-R619W variant either phenocopies the complete loss of PEP or distinctly alters IFN-I production following RLR activation, we cultivated PEP-WT, PEP-null CRISPR, and PEP-R619W BMMs and then stimulated them with LPS or infected with the murine coronavirus Murine Hepatitis Virus Strain A59 (MHV A59), a MDA5 agonist ^27^. There were no differences detected in IFNβ production between PEP-null CRISPR and PEP-WT BMMs following LPS exposure or MHV A59 infection (Figure 2A). There were also no difference detected in IFNβ production between PEP-WT and PEP-R619W BMMs following MHV A59 infection (Figure 2B) or LPS exposure (Figure 2C). These results suggest that PEP does not promote IFN-I production in BMMs following TLR4 or MDA5 activation.

**Figure 2.**
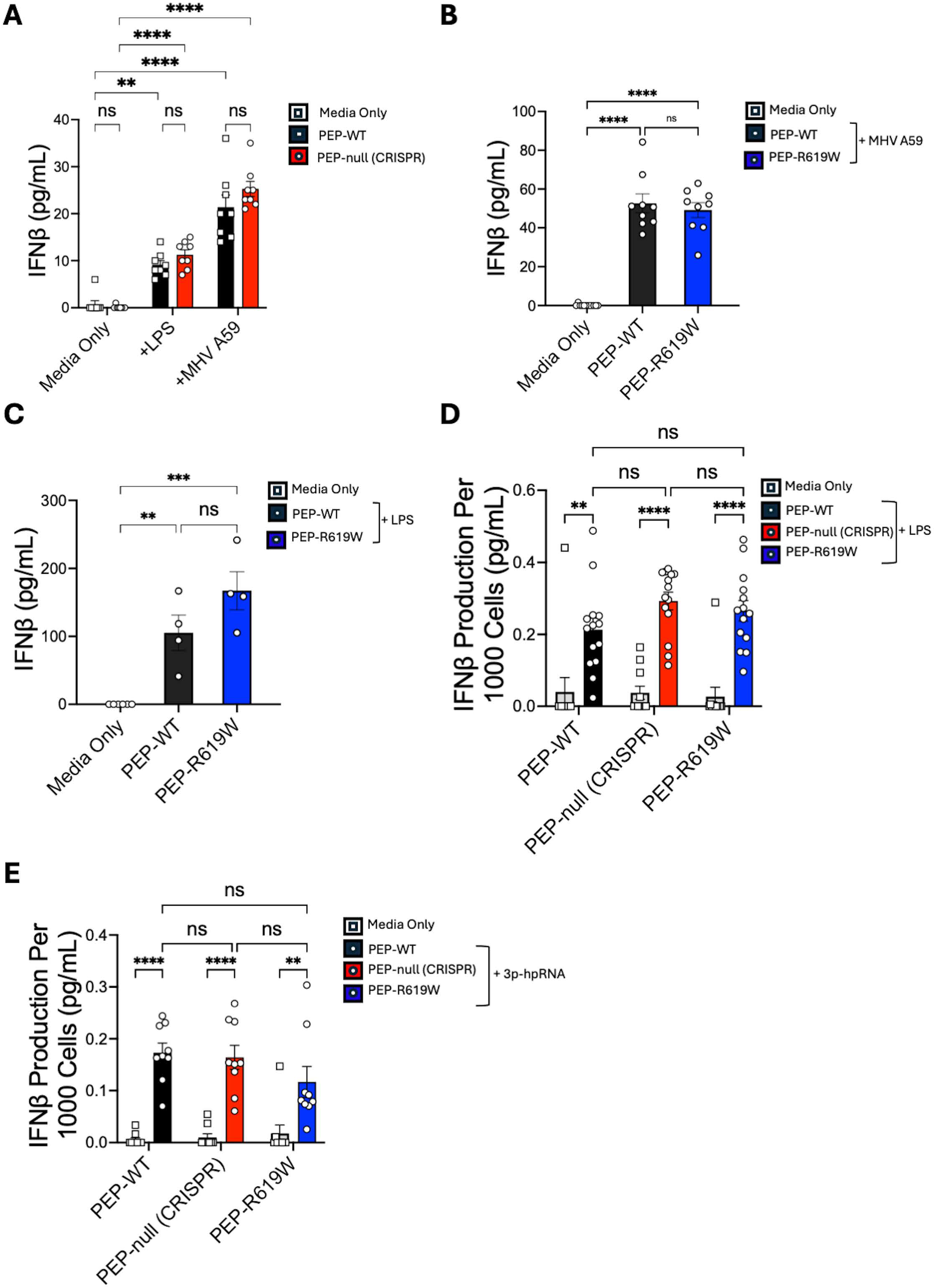
PEP and PEP-R619W do not mediate IFNβ production in myeloid cells following LPS, MHV A59, or viral RNA stimulation. C57BL/6J PEP-WT (black, square symbol), CRISPR-Cas9 generated PEP-null (red, circle symbol), BMMs were cultivated and then stimulated with LPS (500 ng/mL) or infected with MHV A59 (MOI 0.1) for 18 hours **(A)**.C57BL/6J PEP-WT (Black, circle symbol) and CRISPR-Cas9 generated PEP-R619W (Blue, circle symbol) BMMs are cultivated and then stimulated with MHV-A59 (MOI 0.1) **(B)** or with LPS (500 ng/mL) **(C)** for 18 hours. FLT3-L BMDCs were cultivated from C57BL/6J PEP-WT, PEP-null, and PEP-R619W mice. Cultures were incubated with LPS (500 ng/mL) **(D)** or 3p-hpRNA (2 ng/mL) **(E)** for 18 hours. The supernatant was used to determine IFNβ concentration (ELISA). ns= no significance; a Two Way ANOVA with Tukey post hoc analysis was conducted for (A,D,E), a One Way ANOVA was conducted for (B, C). Data is representative of 2 independent experiments **(A)** or pooled from 3 independent experiments **(B-E)**. Each dot represent a biological replicate.

Next, we generated FLT3-ligand (FLT3-L) BMDCs from C57BL/6J PEP-WT, PEP-null CRISPR, and PEP-R619W mice and stimulated them with LPS or 5’ Triphosphate hairpin ribonucleic acid (3p-hpRNA), a RIG-I agonist. We detected no significant differences in IFNβ production between all three genotypes (PEP-WT, PEP-null CRISPR, and PEP-R619W) following either LPS or 3p-hpRNA stimulation (Figure 2 D-E). Taken together, these results indicate that neither PEP nor the PEP-R619W variant modulate IFNβ production in dendritic cells mediated by TLR4 or RIG-I activation.

### PEP and the PEP-R619W variant distinctly modulate the production of pro-inflammatory cytokines in dendritic cells

Our findings show that neither PEP nor the PEP-R619W variant impact IFN-I production in DCs and macrophages during TLR4, MDA5, or RIG-I stimulation. However, DCs produce a variety of other cytokines which can promote widespread inflammation or influence T cell activation and differentiation. Prior studies show that PEP-null and the PEP-R619W variant enhances T cell function through both T cell intrinsic and T cell extrinsic mechanisms during virus infection ^19 28, 29^. Previous studies that have investigated the role of PEP and its variant on cytokine production in myeloid cells have reported varied findings^21, 30, 31^. Using PEP-R619W knock-in mouse models, one group reported increased IL-12 production in Granulocyte-macrophage colony stimulating factor (GM-CSF) differentiated PEP-R619W BMDCs post LPS exposure^30^. Another group reported that GM-CSF differentiated PEP-null JAX BMDCs exhibited enhanced secretion of IL-6, IL-8 and TNFα compared to PEP-WT BMDCs following treatment with muramyl dipeptide (MDP)^31^. They also reported that the loss of PEP reduced while the PEP-R619W variant increased IL-1β secretion by BMDCs following ultra-pure LPS exposure^32^. However, a different group later reported no differences in TNFα, IL-1β, or IL-12 production following treatment with LPS between FLT3-ligand differentiated PEP-WT and PEP-null JAX BMDCs ^21^. These contrary results align with previous reports of PEP-null and PEP-R619W mice exhibiting different phenotypes but could also be attributed to the usage of different mouse models, type of stimulus, and selected assays. Thus, using our CRISPR-Cas9-generated mouse models, we wanted to know if PEP and the PEP-R619W variant regulated the production of other inflammatory cytokines relevant to modulating T cell function.

To accomplish this, we again used FLT3-ligand differentiated BMDCs from PEP-WT, PEP-null CRISPR, and PEP-R619W mice. BMDCs were stimulated with LPS, and the supernatant was collected and analyzed for TNFα, IL-12 (p70), and IL-2 production. We found that compared to PEP-WT BMDCs, TNFα production was reduced in both PEP-null CRISPR and PEP-R619W BMDCs (Figure 3A). Additionally, we observed a reduction in IL-12 production by PEP-R619W BMDCs compared to PEP-WT. However, there was no difference detected in IL-12 production between PEP-null CRISPR and PEP-WT BMDCs (Figure 3B). Finally, we observed increased IL-2 production by both PEP-null CRISPR and PEP-R619W BMDCs compared to PEP-WT BMDCs (Figure 3C). Taken together, these findings suggest that PEP and the PEP-R619W variant distinctly modulate cytokine production in dendritic cells.

**Figure 3.**
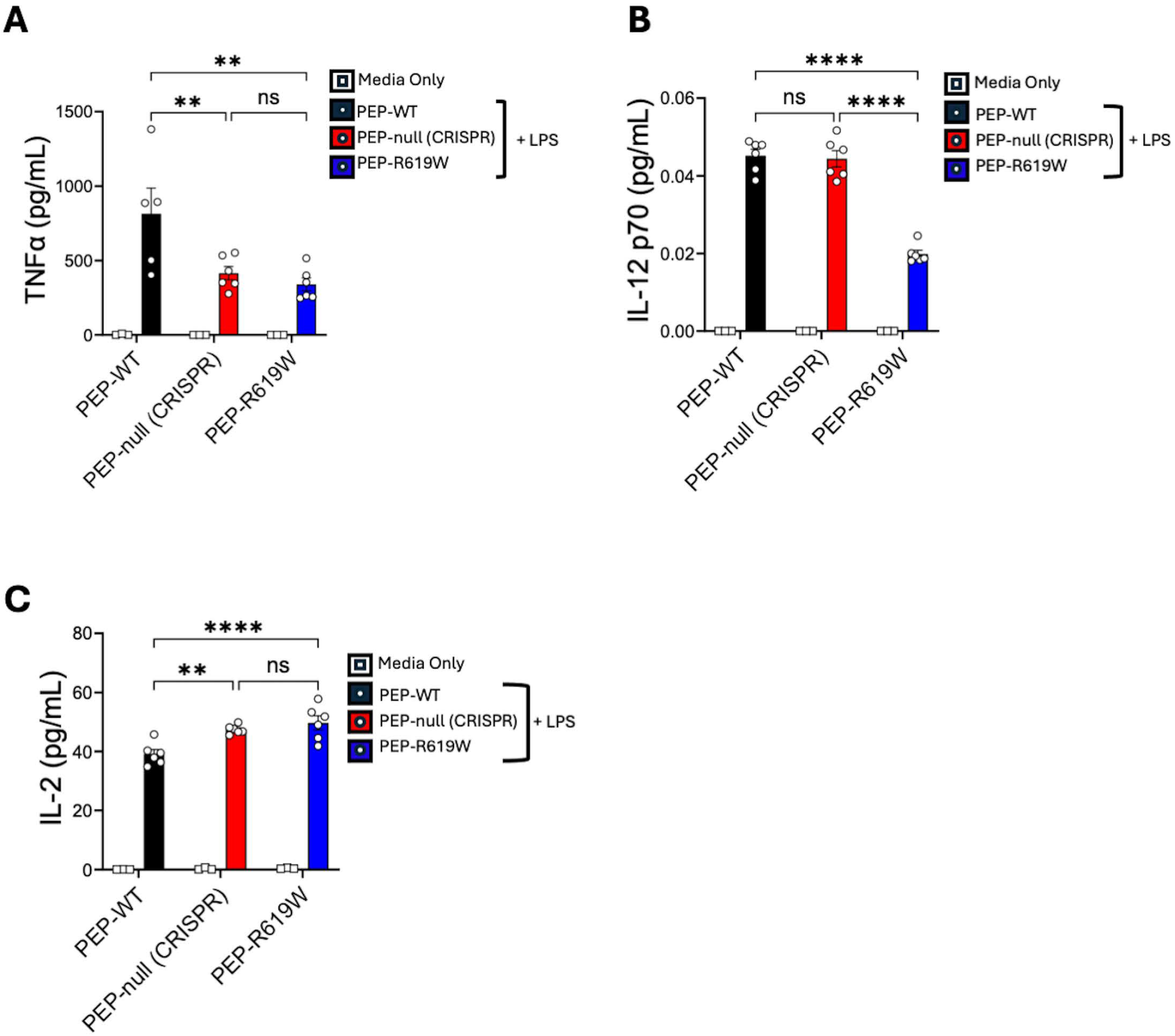
PEP and the PEP-R619W variant distinctly modulate the production of pro-inflammatory cytokines in dendritic cells. FLT3-L BMDCs were cultivated from C57BL/6J PEP-WT (black), CRISPR-Cas9 generated PEP-null (red), and PEP-R619W (blue) mice. Cultures were incubated with LPS (500 ng/mL) for 18 hours. The supernatant was used to determine TNFα **(A)**, IL-12 **(B)**, and IL-2 **(C)** concentration (ELISA). ns= no significance;**p<0.01, ****p<0.0001; Two Way ANOVA with Tukey post hoc analysis. Data is representative of 2 independent experiments. Each dot represents a biological replicate.

## DISCUSSION

The *PTPN22* 1858C>T allele is strongly associated with the development of autoimmunity, but the murine equivalent mutation has also been shown to be protective during virus infection and cancer^18, 19, 33^. While the impact of the PEP-R619W variant is well-defined in T cells, the molecular mechanisms by which PEP and its variant impact the function of other immune cells, such as the ability of APCs to produce pro-inflammatory cytokines, remain less understood. Previously, it was suggested that PEP promotes IFN-I production in macrophages and dendritic cells^21^. However, these data were generated using PEP-null mice on a C57BL/6J;C57BL/6N mixed genetic background, posing the concern that our current understanding of PEP’s role during IFN-I production may have been influenced by the C57BL/6N SNPs present in the PEP-null JAX mouse model. Using our CRISPR-Cas9-generated PEP-null model, we first set out to determine if previous data regarding the impact of PEP on IFN-I production could be recapitulated. While the PEP-null JAX BMDCs had reduced IFN-I production, we did not detect a difference between our PEP-null CRISPR BMDCs and C57BL/6J PEP-WT BMDCs (Figure 1B). These strain-specific differences in IFN-I production recapitulate previous findings, while also revealing that strain specific difference, likely genetic variation between C57LB/6J and C57Bl/6N mice, in the PEP-null JAX mice may have influenced the previously observed phenotype. This interpretation is further supported by reports of reduced inflammatory responses during virus infection in C57BL/6N mice compared to C57BL/6J mice ^34, 35^. However, our study results do not address why PEP-null JAX BMMs engineered to overexpress the human protein Lyp have restored IFNβ production ^21^. Identifying the mechanism that drives the reduced IFNβ production observed in the C57BL/6J;C57BL/6N PEP-null JAX mice but not in the PEP-null CRISPR mice will require further investigation. Regardless, we are not the first to comment on or identify discrepancies regarding previous data reported on the role of PEP during IFN-I production in myeloid cells. However, these prior publications have largely focused on differences identified in human samples, as well as in the context of systemic lupus erythematous (SLE) ^36-40^. Thus, to the best of our knowledge, we are the first to show in a mouse model that there may be strain-specific differences confounding the interpretation of how/if PEP regulates cytokine production in DCs and macrophages.

Next, when evaluating the specificity of the PEP inhibitor LTV-1 during IFN-I production in BMMs, we found reduced IFNβ production in both PEP-WT and PEP-null CRISPR macrophages treated with LTV-1 (Figure 1C). These findings suggest that the observed reduction of IFNβ may not be due to the deficiency of PEP nor the inhibition of its enzymatic function, but rather because of non-specific effects of LTV-1. Taken together, our findings reveal additional confounding variables that could impact the results of studies that aim to discern the role of PEP during inflammation through the use of either genetic models or chemical inhibition. Our observations add to the numerous conflicting findings already identified in the field regarding PEP, PEP-R619W, and their effects on cytokine production ^21, 30-32^. These differences likely stem from genetic variation present in both human and animal studies, as well as the utilization of diverse experimental approaches including over-expression, transgenic models, knockdown techniques, and chemical inhibition. Each of these systems offer distinct advantages yet also varying limitations, yielding inconsistent results. To address this lack of clarity, our lab employs CRISPR/Cas9-generated PEP-null and PEP-R619W mouse models on a C57BL/6J background. By using this model, we can precisely evaluate the mechanisms by which PEP and the PEP-R619W variant drive inflammation while minimizing confounding genetic variables.

Using our PEP-null CRISPR and PEP-R619W mice, our results indicate that IFNβ production by DCs and macrophages is not promoted by PEP or the PEP-R619W variant following TLR4, MDA-5, or RIG-I stimuli (Figure 1B, Figure 2A-C). Although our findings indicate that PEP and PEP-R619W do not promote IFNβ production in myeloid cells, we found that PEP and the PEP-R619W variant distinctly modulate the production of other pro-inflammatory cytokines. More specifically, we found that compared to PEP-WT, TNFα production was reduced in both PEP-null CRISPR and PEP-R619W DCs (Figure 3A). Additionally, we observed a reduction in IL-12 production by PEP-R619W DCs compared to PEP-WT but no significant change in the production of IL-12 was detected in the PEP-null CRISPR BMDCs when compared to PEP-WT BMDCs (Figure 3B). Previously it was reported that PEP-R619W GM-CSF differentiated BMDCs have increased IL-12 production compared to PEP-WT ^30^. These conflicting findings may be due to a variety of factors such as differences in the PEP-R619W mouse models, LPS dosage, and/or functional differences between FLT3-ligand differentiated and GM-CSF differentiated BMDCs. Finally, we observed increased IL-2 production by both PEP-null CRISPR and PEP-R619W BMDCs compared to PEP-WT. Since DC derived IL-2 can stimulate the activation of T cells, it is possible that the increased IL-2 produced by PEP-null CRISPR and PEP-R619W DCs may be a contributing factor to the enhanced T cell function observed during virus infection ^19, 28, 41^.

Altogether, our findings challenge the current understanding of PEP while providing new insight into how the PEP-R619W variant is capable of simultaneously promoting autoimmunity and conferring protection against persistent viral infection and cancer. While our results reveal that changes in IFN-I production are not a key factor in how PEP regulates inflammation as previously regarded, our data indicates that PEP and the PEP-R619W variant have a novel role in modulating the production of other cytokines by dendritic cells which are critical for a variety of inflammatory processes. Future studies will need to evaluate how changes mediated by PEP and the PEP-R619W variant in dendritic cells and other antigen presenting cells are relevant in order to accurately define the broad consequences they have on disease pathogenesis.

## MATERIAL AND METHODS

### Ethics

All animal studies were reviewed and approved by University of Kansas Animal Care and Use Committee (IACUC) (protocol number: 278-01).

### Mice

Mice were bred and housed in specific pathogen free general housing conditions at University of Kansas (Lawrence, KS). All animal studies were reviewed and approved by the University of Kansas Animal Care and Use Committee (protocol number: 278-01). Both males and females ranging from 5-12 weeks of age were used in this study. C57BL/6J WT (PEP-WT) mice were originally purchased from Jackson labs, then bred and maintained in the University of Kansas Animal Care Unit. Ptpn22 ^-/-^ (PEP-null CRISPR) mice were generated using CRISPR/Cas9 technology on a C57BL/6J background, as described previously, by Dr. Kerri Mowen and Dr. Linda Sherman (Scripps Research Institute, La Jolla, CA) and were gifted from Dr. Sherman^42^. Mice expressing the Ptpn22 pro-autoimmune allele (PEP-R619W) were generated using CRISPR/Cas9 technology on a C57BL/6J background using methods previously reported by Dr. Kerri Mowen and Dr. Linda Sherman (Scripps Research Institute, La Jolla, CA) and were gifted from Dr. Sherman^18, 19, 42^. Approximately every 10 generations, PEP-null CRISPR and PEP-R619W mice are backcrossed to new C57BL/6J wildtype mice to reduce the likelihood strain specific SNPs have arisen and driving a particular phenotype. Mice from the same parental pairs are not used in breeding pairs. B6.Cg-Ptpn22^tm2Achn^/J Ptpn22 ^-/-^ mice (PEP-null JAX) were gifted from Dr. Won Jin Ho (John Hopkins University School of Medicine, Baltimore, MD). These mice are available for purchase from the Jackson Laboratory (strain number #028977), and their generation has been previously described^22^.

### Bone Marrow derived Macrophage (BMM) Culture

To generate BMMs, bone marrow cells were isolated from PEP-WT, PEP-null JAX, PEP-null CRISPR and PEP-R619W mouse femurs. Following isolation, cells were cultured in Roswell Park Memorial Institute medium (RPMI) supplemented with 10% FBS, 1% L-glutamine, and 1% Penicillin/Streptomycin with 50ng/mL M-CSF (Stem Cell Technologies, Vancouver, CA) for 8 days. An additional 10 mL of M-CSF-containing media was added to BMM cultures on day 3, and on day 8, differentiated cells were harvested, counted, and replated for further assays.

### Bone Marrow Dendritic Cell (BMDC) Culture

Bone marrow cells were isolated from PEP-WT, PEP-null, and PEP-R619W mouse femurs. Following isolation, bone marrow cells were cultured in Advanced DMEM containing 10% FBS, 1% penicillin streptomycin, 1% L-Glutamine, and 100ng/mL FLT3L (Stem Cell Technologies) for 8 days. On day 8, FLT3-ligand differentiated dendritic cells were harvested, counted, and re-plated for further assays.

### Myeloid cell Stimulation

#### BMM stimulation

Bone marrow macrophages (BMMs) were plated in a 96-well plate at a density of 5 x 10^4^ cells per well. For infection, cells were inoculated with MHV-A59 at a multiplicity of infection (MOI) of 0.1. The plate was incubated for one hour at 37°C with shaking every 10 minutes to ensure even distribution of the virus. Following the one-hour incubation, the plate was centrifuged at 300g for 10 minutes to pellet the cells. The supernatant containing the unadsorbed virus was carefully discarded. The cells were then resuspended in fresh media and incubated at 37°C for 18 hours to allow for viral replication. After the 18-hour incubation, the cell culture supernatant was collected and stored at -80°C for subsequent IFNβ ELISA.

#### BMDC stimulation

BMDCs were resuspended and plated at 1 x 10^6^ cells/mL in Advanced DMEM (Gibco) containing 10% FBS, 1% penicillin streptomycin and 1% L-Glutamine. The cells were stimulated with either LPS or 3p-hpRNA. Cells stimulated with LPS (500 ng/mL) (Sigma) were incubated overnight (16-18 hours). The supernatant was collected and stored frozen at -80 degrees Celsius for future assays. Cells stimulated with 3p-hpRNA (2 ng/mL) complexed with LyoVec (InvivoGen) were incubated overnight (16-18 hours). The supernatant was collected and stored frozen at -80 degrees Celsius for future assays.

### Enzyme-Linked Immunosorbent Assay (ELISA)

Levels of IFNβ, TNFα, IL-12 (p70), and IL-2 in cell culture supernatant were measured using ELISA Kits (IFNβ (PBL Assay Science), TNFα, IL-12 and IL-2 (BioLegend)) according to the manufacturer’s instructions, followed by analysis with a BioTek EON microplate reader at 450 and/or 570 nm.

### qRT-PCR

RNA was extracted from 6-8 week old PEP-WT and PEP-R619W mouse splenocytes using TRIzol according to the manufacturer’s instructions (Invitrogen). cDNA was synthesized using the High-Capacity cDNA Reverse Transcription Kit (Applied Biosystems), according to the manufacturer’s instructions. Real-time PCR was performed on a Quant Studio 3 system using PowerTrack^TM^ SYBR^TM^ Green Master Mix (Applied Biosystems). Each reaction was measured in duplicate, and data was normalized to the expression levels of the house keeping gene *Gapdh*. Primers for the genes were as follows: *Gapdh* Forward Primer 5’ ACGACCCCTTCATTGACCTC 3’ and Reverse Primer 5’ ACTGTGCCGTTGAATTTGCC 3’; *Ptpn22* Forward primer 5’ AGCTGATGAAAATGTCCTATTGTGA 3’ and Reverse primer 5’ GTCCCACTGCATTCTGGTGA 3’.

### Statistical analysis and graphing

All statistical analysis was performed using GraphPad Prism Software. The type of statistical test is listed in figure legends. Data was considered statistically significant if the p value<0.05. Graphs were made in GraphPad Prism (La Jolla, CA). Figure legends indicate if data shown is pooled from multiple studies or is from representative study.

## ACKNOWLEDGEMENTS

The PEP-R619W and PEP-null CRISPR mice were originally generated by Dr. Kerri Mowen and Dr. Linda Sherman, Scripps Research Institute. These mice were gifts from Dr. Sherman. PEP-null JAX mice were a gift from Dr. Won Jin Ho (Johns Hopkins University School of Medicine). MHV A59 was a gift from Dr. Anthony Fehr (University of Kansas).

## AUTHOR CONTRIBUTIONS

Jenna R. Barnes: Data Curation, Formal Analysis, Investigation, Methodology, validation,

Visualization, Writing-original draft, writing-review and editing

Anam Fatima Shaikh: Data Curation, Formal Analysis, Investigation, Methodology, Validation,

Visualization, Writing-original draft, writing-review and editing

Alec M. Bevis: Data Curation, Formal Analysis, Investigation, Methodology, Validation,

Writing-original draft, writing-review and editing

Tammy R. Cockerham: Methodology, Investigation, Project Administration

Robin C. Orozco: Conceptualization, Formal Analysis, Funding acquisition, Project administration, Supervision, Validation, Visualization, Writing-Original draft, Writing-review and edit

## DATA AVAILABILITY

All data necessary to interpret results is presented in this manuscript. Raw data will be made available upon request.

## ABBREVIATIONS

IFN-I: Type I Interferon
Ptpn22: Protein tyrosine phosphatase non-receptor type 22
Lyp: Lymphoid protein
PEP: PEST-domain enriched phosphatase
DCs: Dendritic Cells
TLR4: Toll-Like Receptor 4
LPS: Lipopolysaccharide
3p-hp-RNA: 5’ Triphosphate hairpin ribonucleic acid
MHV A59: Mouse Hepatitis virus strain A59
TNFα: Tumor Necrosis Factor alpha
IL: Interleukin
R: Arginine
W: Tryptophan
HLA: Human Leukocyte Antigen
Tregs: T regulatory cell
CRISPR-Cas9: Cluster regularly interspaced short palindromic repeats Cas9
APC: Antigen Presenting Cell
LCMV-cl13: Lymphocytic Choriomeningitis Virus Clone 13
TLR3: Toll-like receptor 3
SNP: Single Nucleotide Polymorphism
RLR: Rig I like receptor
MDA5: Melanoma Differentiation Associated protein 5
IFNβ: Interferon Beta
RNA: Ribonucleic Acid
BMDC: Bone Marrow Derived Dendritic cells
BMM: Bone Marrow Derived Macrophage
JAX: Jackson Labs
mRNA: Messenger Ribonucleic Acid
FLT3-L: FMS-like tyrosine kinase 3-Ligand
GM-CSF: Granulocyte-Macrophage Colony Stimulating factor
RIG-I: Retinoic acid inducible gene 1
MDP: Muramyl dipeptide
ELISA: Enzyme Linked Immunosorbent Assay
ANOVA: Analysis of Variance
NFκB: Nuclear Factor kappa B
TRAF3: TNF receptor-associated factor 3
IACUC: Institutional Animal Care and Use Committee
DMEM: Dulbecco’s Modified Eagle Medium
FBS: Fetal Bovine Serum
MOI: Multiplicity of Infection
Ns: Not Significant
qRT-PCR: Quantitative Reverse Transcriptase Polymerase Chain Reaction

